# Cell segmentation using deep learning: comparing label and label-free approaches using hyper-labeled image stacks

**DOI:** 10.1101/2020.01.09.900605

**Authors:** William D. Cameron, Alex M. Bennett, Cindy V. Bui, Huntley H. Chang, Jonathan V. Rocheleau

## Abstract

Deep learning provides an opportunity to automatically segment and extract cellular features from high-throughput microscopy images. Many labeling strategies have been developed for this purpose, ranging from the use of fluorescent markers to label-free approaches. However, differences in the channels available to each respective training dataset make it difficult to directly compare the effectiveness of these strategies across studies. Here we explore training models using subimage stacks composed of channels sampled from larger, ‘hyper-labeled’, image stacks. This allows us to directly compare a variety of labeling strategies and training approaches on identical cells. This approach revealed that fluorescence-based strategies generally provide higher segmentation accuracies but were less accurate than label-free models when labeling was inconsistent. The relative strengths of label and label-free techniques could be combined through the use of merging fluorescence channels and out-of-focus brightfield images. Beyond comparing labeling strategies, using subimage stacks for training was also found to provide a method of simulating a wide range of labeling conditions, increasing the ability of the final model to accommodate a greater range of experimental setups.

## I Introduction

As high-content, high-spatiotemporal cellular imaging becomes more widespread, the ability to perform cellular segmentation both quickly and accurately becomes increasingly critical for efficient cellular analysis and feature extraction. Advances in deep learning have positioned neural networks as a powerful alternative to traditional approaches such as manual and algorithmic-based segmentation [1], [2]. In particular, the development of the U-Net architecture provided a significant boost to segmentation performance [3], [4] and has now become the template for many modern segmentation models [5], [6], [7]. Advancements in our understanding of deep learning have also made the technique more accessible for smaller-scale operations. Techniques such as data augmentation have significantly reduced dataset size requirements, while improvements to training (e.g. transfer learning, initialization, dropout, hyperparameter schedulers, and optimizers) have reduced training times considerably [8], [9], [10], [11].

Adapting deep learning techniques for cellular segmentation, however, presents some unique challenges due to the high levels of incongruity across images of cells. For instance, images of cells may appear significantly different as a result of either the imaging techniques [12], or even the imaging settings used (e.g. using a different numerical aperture or magnification of the objective lens). Furthermore, the cells themselves may demonstrate an enormous range of morphologies depending on factors such as cell type, cell confluency, and local environment. Cell appearance may also vary significantly depending on the cellular structures targeted through the experimental labels or dyes used. To address this variety, many segmentation algorithms are highly-tailored to their target application and therefore do not experience widespread adoption [13]. The specialized nature of each dataset and resulting solution also means that it is difficult to compare individual labeling approaches or segmentation strategies and establish best practices. The challenges when developing a cell segmentation approach for novel applications are therefore three-fold: (i) developing a method of assessing which imaging or labeling strategies produce the greatest segmentation accuracies, (ii) developing strategies to efficiently train models capable of maintaining an acceptable level of segmentation accuracy across a wide range of potential input configurations, and (iii) keeping dataset requirements small enough that they are feasible to retrain for new applications when necessary.

Traditional approaches to automated cell segmentation from microscope images generally fall into two main categories: fluorescence-based and label-free approaches. Fluorescence-based approaches often boast higher segmentation accuracies but require the addition of fluorescence markers that can also induce stress on the cell, either directly or as a byproduct of imaging and are therefore best avoided when possible [14], [15], [16]. For genetically-encoded sensors, the successful co-expression of the desired sensors and markers in a single cell becomes increasingly difficult in hard-to-transfect cell lines, which limits the population of cells that can be both successfully segmented and analyzed. Reliance on specific fluorescence markers can also confer some significant disadvantages as microscopy trends towards multiparametric, high-throughput imaging. Most notably, fluorescence-based segmentation limits multiparametric imaging by dedicating a portion of the fluorescence spectra for segmentation that might otherwise be used. In contrast, label-free approaches (e.g. brightfield imaging) have the advantage of not requiring a fluorescent marker, but often struggle with reduced performance in high confluence when the boundaries between cells are not distinct [3].

Many experiments use fluorescence sensors or dyes for reasons extraneous to segmentation (e.g. as a source of data collection and localization), which represents a valuable opportunity to augment the data presented to a segmentation model and thereby improve performance. For instance, genetically-encoded sensors expressed in either the cytoplasm or the mitochondria may each be used to help demarcate and mask individual cells. However, the information provided by these signals are very different: cytoplasmic markers can clearly define cell boundaries in isolation, but may become indistinct across adjacent cells; mitochondrial markers do not reach the limits of cell boundaries, but provide a gap in fluorescence which can be used to more broadly separate adjacent cells. The primary challenge when using this information is that the combination of auxiliary fluorescence signals available to the segmentation model may vary from experiment to experiment or cell to cell. This requires that a segmentation model be trained to maintain performance across a wide range of potential experimental labeling conditions, including the absence of fluorescence labels of any kind. This would be difficult under the traditional approach to training deep learning models, as accounting for all possible experimental configurations would require collecting and labeling a prohibitively large and expensive dataset. Fortunately, microscope images possess a unique property that may be exploited to substantially reduce this dataset requirement: the channels of microscope images exist as stacks of independent images. This means that a subset of channels from a larger image stack can be assembled to create an entirely new, representative microscope image. For example, an image stack of a cell composed of channels capturing a cytoplasmic marker, a membrane marker, a mitochondrial marker, and a brightfield image can be used to simulate a cellular image where only the cytoplasmic marker is present. The ability to create representative image substacks from a larger image stack is in stark contrast to other vision-based image modalities (e.g. object detection using camera footage), where removing a specific color channel would produce an image that is no longer representative of the target data. Here, we use source image stacks composed of three fluorescent labels and brightfield images imaged at seven different focal planes to simulate a wide range of expected experimental images (see **Fig. 1** and **Supplementary Fig. S2**) to train a robust cell segmentation model. For simplicity, we henceforth refer to the source image stack as the **hyper-labeled image stack** and subsets of the images used during training as **subimage stacks**. Beyond reducing the dataset size, this approach to training confers some additional advantages, including the ability to compare different labeling approaches on an identical dataset. In particular, we use this dataset to explore new approaches to pre-process data entering segmentation models including the use of out-of-focus (OoF) brightfield images and the concept of merging fluorescence channels into a single input channel. To keep the implementation practical, our approach uses fewer than 300 labeled cell examples and can be trained in less than a day on a modern GPU.

**Fig. 1.**
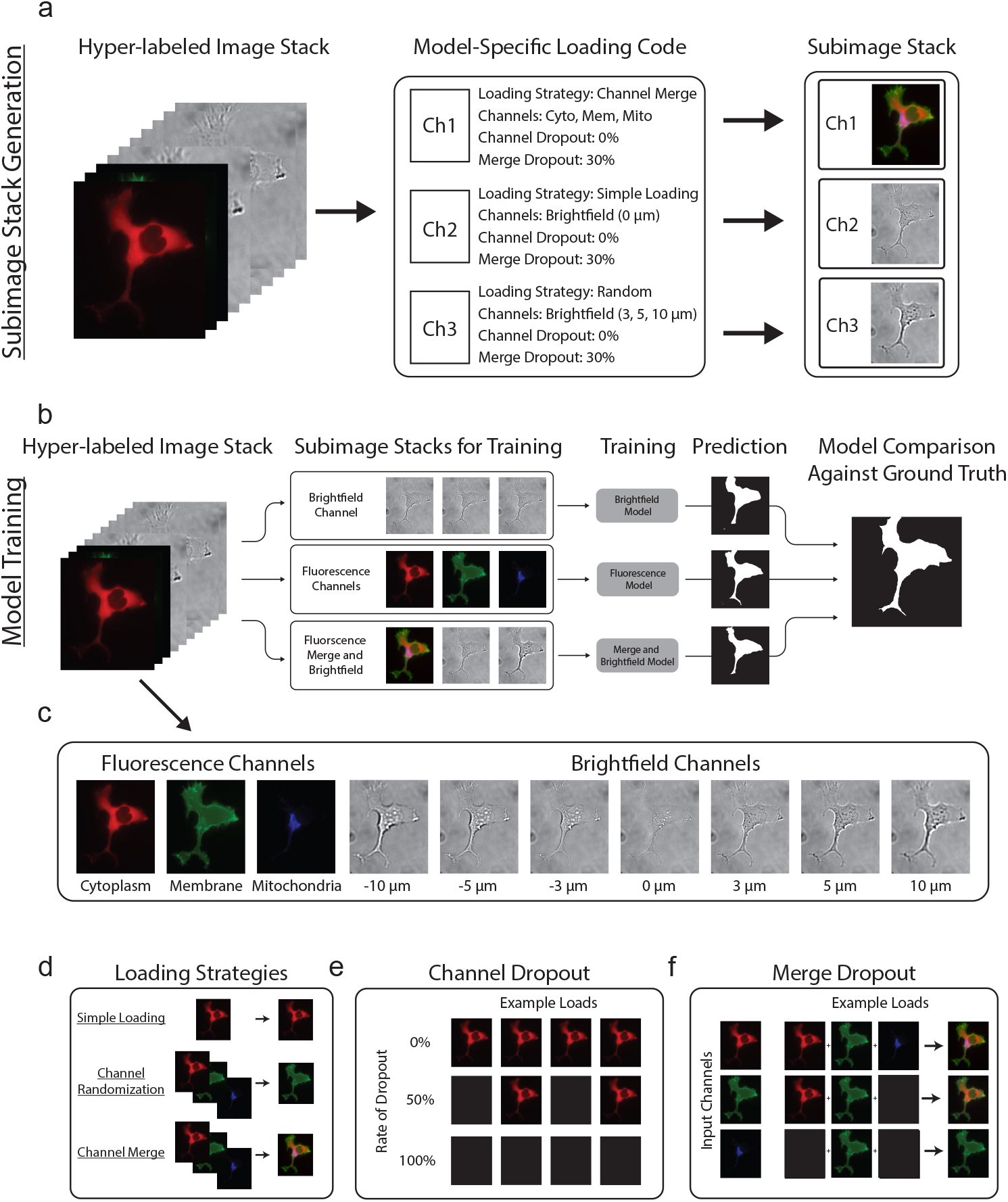
Overview of the dataloader. (a) Three-channel subimage stacks are assembled from a source, or hyper-labeled, image stack according to a model-specific loading code. (b,c) Segmentation models are trained independently using their unique subimage stacks. Each model begins with the same pretrained model and the same training-validation split for a given replicate. Once training is complete, model accuracy is assessed by comparing the average accuracy of each model over the final 10 training epochs. Note: the prediction image shown is from the example hyper-labeled image for simplicity; however, the actual validation is composed of images not seen during training. (d) Individual channels are constructed using three core techniques: i) Simple loading, which loads the channel directly, ii) Randomization, which chooses a channel at random from a preset subset of channels, iii) Channel merging, which averages the intensities from multiple channels into a single channel. (e) Channels may also be affected by a “channel dropout” rate, which determines the probability that an individual channel will be replaced by a blank channel. (f) Channel merge may also be affected by its own “merge dropout” rate, which determines the probability that a blank channel will be substituted in place of that particular channel during the merge.

## II Methods

### A. Dataset Collection

Cells were labeled with three distinct subcellular markers for the cytoplasm (mTurquoise2-tagged Apollo-NADP^+^) [17]), the membrane (YFP-Mem), and the mitochondrial matrix (Mitotracker Deep-Red FM). Imaging was performed using a three-colour widefield fluorescence microscope from ASI. Additionally, brightfield images were taken of each cell at 7 different focal depths (−10, −5, −3, 0, +3, +5, +10 μm), with 0 μm representing cells in manual focus. Together, these 10-channels were used to form our hyper-labeled image stacks. From these images, 275 fully labeled cells (cells containing all three fluorescent markers) were manually isolated using a custom Fiji plugin to form our training and validation datasets. Ground-truth segmentation labels were created manually by alternating between the fluorescence and near-focus (−3 to 3 μm) brightfield channels to minimize biasing performance towards either imaging method. This final dataset represents a combination of 106 AD293 and 169 INS1E cells, with an additional 175 cell-free images added to the dataset as negative controls. Of the cellular images, approximately 26% represented isolated cells (i.e. no direct contact with other cells) while the remaining 74% had at least one neighboring cell in the field of view. Cell images were resized to a final size of 400 × 400 px before entering the model.

An additional test set was created as described above using cells that were labeled with subcellular markers for the endoplasmic reticulum (mTurquoise2-tagged Apollo-NADP+), the membrane (YFP-Mem), and the mitochondrial matrix (Mitotracker Deep-Red FM). This dataset was composed of 96 fully labeled cell images and 60 cell-free negative controls. The overall methodology was confirmed on an externally sourced dataset [18]. Briefly, the macrophage cells in the dataset were labeled with two distinct subcellular markers for the cytoplasm (BODIPY 493/503) and the nucleus (Sytox). Image stacks of these cells were composed of both fluorescence channels with additional brightfield images taken at 5 different focal depths. This dataset represents a combination of 264 labeled cells and an additional 122 cell-free images as negative controls.

### B. Model and Training Parameters

Training was performed on a variation of the U-Net model [3], which employs a descending arc (contracting path) to increase feature information followed by an ascending arc (expansion path) to combine feature and spatial information. The model used here was composed of a traditional ResNet34 [19] architecture for the descending arc path and a custom ascending arc that used pixel shuffling during upsampling to reduce checkerboard artifacts [20] (see **Supplementary Fig. S1**). To make use of transfer learning, pretrained weights (provided by [21]) from the ResNet34 model were used for the descending arc while the ascending arc was randomly initialized.

The majority of the hyperparameters used represent best-practice recommendations as described in [21]. However, the learning rate, number of training epochs, and cross entropy weights were determined experimentally. Learning rate was scheduled as a variation of the 1cycle policy [11], [21] (**Supplementary Fig. S3a,b**). The maximum learning rate was chosen by training the model over 100 iterations while gradually increasing the learning rate from 1 × 10^−7^ to 1 × 10^−1^ and recording the training loss. The learning rate chosen was found in the area of the steepest downward slope before loss started to rapidly increase for any of the configurations tested (**Supplementary Fig. S3c,d**). As training was performed in two parts (5 epochs with the pre-trained descending arc frozen, then 100 epochs with the entire network unfrozen), two maximum learning rates were chosen. Based on these results, the maximum learning rate was chosen conservatively as 2 × 10^−4^ for the first 5 epochs, then increased from 2 × 10^−6^ to 1 × 10^−4^ across the network’s parameter groups for the final 100 epochs.

Models were trained using a two-class weighted cross entropy loss:

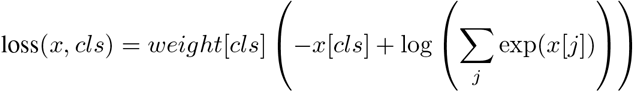

where *weight*[*cls*] and *x*[*cls*] are the weight and prediction for a particular class, *cls*, and *x*[*j*] are the predictions for each individual class (cell or background). The use of crossentropy loss provides the opportunity to weigh the losses related to the cell or the background differently. Altering the background weight with respect to the cell weight produced conservative segmentations at low values and overeager segmentations at high values (**Supplementary Fig. S3e**). Overall accuracy declined in either direction; however, lower values were associated with fewer background pixels being misclassified as belonging to the cell, while the inverse was true for larger weights (**Supplementary Fig. S4**). As many cell masking applications would prefer falling slightly short of the cell boundary (false negative pixels) over surpassing the cell boundary (false positive pixels), a scaling factor of 0.5 was chosen to minimize false positive pixels without significantly impacting overall accuracy.

Training and validation sets were divided using a randomized 80:20 split of the cells from the complete dataset. To determine how the split between the training and validation set may impact model accuracy, models were trained on all-black images using either a consistent (100 models) or a re-randomized (100 models) training-validation split (**Supplementary Fig. S3f,g**). Significant, normally-distributed variability was found in the final training accuracy when the dataset was randomly split. When the dataset was consistently split, the accuracy did not vary. This indicates that it is critical to use identically-split training and validation datasets across each model tested during a single experimental replicate. This also suggests that the relative performance of each model is a more useful metric than an absolute segmentation accuracy percentage. Based on these findings, splits were kept consistent across models during a run to reduce the impact of training-validation splits on relative performance.

### C. Data Augmentation

Significant data augmentation was used to keep the dataset relatively small. As outlined in **Supplementary Fig. S5**, examples of brightfield and fluorescence channels were passed through a variety of transforms including: brightness and contrast, dihedral transforms, image flipping, image jitter, perspective warping, image rotation, skew, and symmetric warping. Parameter ranges for each transform were chosen that produced realistic cell images for both fluorescence and brightfield images. Despite the unnatural appearance, zero-padding was used when necessary to avoid twin-cell artifacts in the ground-truth labels (i.e. the presence of two labeled cells). Data-augmentation beyond a squaring crop and resize was not applied during validation.

### D. Subimage Stack Generation

Training was performed on three-channel subimage stacks assembled from the ten-channel, hyper-labeled image stacks. Creation of these three-channel subimage stacks was performed during training using model-specific loading codes. These loading codes provided independent, channel-specific instructions to a custom dataloader directing how each of the three channels would be assembled from the source dataset (see **Fig. 1**). The use of loading codes permitted more complex interactions with the source image stack’s channels, providing the dataloader with the following abilities:

- **Simple Loading**– Load a specific channel from the ten-channel source image stack
- **Randomization**– Load a single random channel from a predefined subset of channels
- **Channel Dropout**– Perform a randomized test against a dropout percentage. Load the channel normally if passed; load a blank channel otherwise
- **Merge**– Merge the contents from multiple channels into a single channel before loading
- **Merge Dropout**– Perform a randomized test against a dropout percentage and only include that particular channel in the merge if passed

A complete representation of the loading combinations used can be found in **Supplementary Fig. S2**.

### E. Plotting and Metrics

Model performance was determined using their average segmentation accuracy, which was calculated as the percentage of pixels that were accurately classified as compared to the ground-truth label (also known as pixel accuracy). Unless otherwise noted, line graphs were plotted using a ten-point moving average for clarity (with empty padding for early epochs) and error bars represent the standard error of the mean. Significance was assessed using a paired t-test.

### F. Availability of Code and Dataset

Training code and links to download the dataset are available through Github at: https://github.com/RocheleauLab

## III Results

### A. Comparing segmentation accuracies when using common cell labeling approaches

Segmentation models were trained using three-channel subimage stacks generated from the ten-channel hyper-labeled image stacks (**Fig. 1a** and **Section II-D**). This approach allows a direct comparison of labeling strategies using a diverse set of source inputs (e.g. label-free segmentation using only brightfield images or fluorescence-based segmentation using a combination of cytoplasmic, membrane and mitochondrial markers; **Fig. 1b,c**). Generation of subimage stacks in this manner also permits more advanced features such as channel randomization, channel merge, channel dropout, and merge dropout (**Fig. 1d-f**). In particular, adding a channel dropout rate can be used to simulate varying expression of a particular fluorescence channel across cells (e.g. cells that are variably labeled with a cytoplasmic tag; **Fig. 1e**). The ability to simulate variable expression is critical for training models where any fluorescence channel may vary across cells, experiments, or even channels. This is particularly true when auxiliary fluorescence signals are used for cell segmentation.

Using this approach, we compared the performance of segmentation models trained on distinct subimage stacks loaded with either: a single fluorescence marker (the cytoplasmic, membrane, and mitochondria models), the in-focus brightfield channel (the brightfield model), a combination of all three fluorescence markers (the fluorescence model), random channels (the random model), or black channels as a negative control (the ‘all black’ model). **Supplementary Fig. S2** provides an example of each subimage stack. Training was performed for 105 epochs, which was sufficient for all models to reach at least 90% of their peak accuracy (**Fig. 2a**). Segmentation performance of the fluorescence, cytoplasmic and membrane models were significantly better than the other approaches (**Fig. 2b**). More generally, single-channel fluorescence images performed well when the fluorescence touched the cell boundary (cytoplasm and membrane, *>*96%), but poorly when this was not the case (mitochondria, reaching ~92%). The brightfield model performed only slightly better than the mitochondrial equivalent (reaching ~93%), but is not influenced by labeling conditions as is the case for the models based on fluorescence markers. To determine how variability in labeling would affect the performance of fluorescence-based models, a channel dropout rate (as outlined in **Fig. 1e**) was added to each fluorescent channel ranging from 0% to 100% with the latter representing black input channels (**Fig. 2c,d**). As the level of dropout increased, performance converged to that of the all-black control in all models with the exception of brightfield. This was particularly devastating for models relying on a single fluorescent marker (cytoplasm, membrane, mitochondria), where brightfield performance began to surpass that of the cytoplasmic or membrane models at ~20% and ~30% dropout, respectively. In contrast, the use of three distinct fluorescent markers in the fluorescence model allowed it to suffer a dropout rate of ~50% before performance dipped below that of the brightfield model. The brightfield model was particularly poor at distinguishing cell boundaries when cells were highly confluent (**Fig. 2e**). These data highlight the value of using specific fluorescent signals to improve segmentation performance as compared to brightfield alone; however, relying exclusively on fluorescence may significantly impact performance when labeling is inconsistent. Furthermore, these data suggest that only certain fluorescent signals offer an improvement over label-free approaches (e.g. membrane and cytoplasmic markers), and that brightfield may be the more effective option in others (e.g. the mitochondria).

**Fig. 2.**
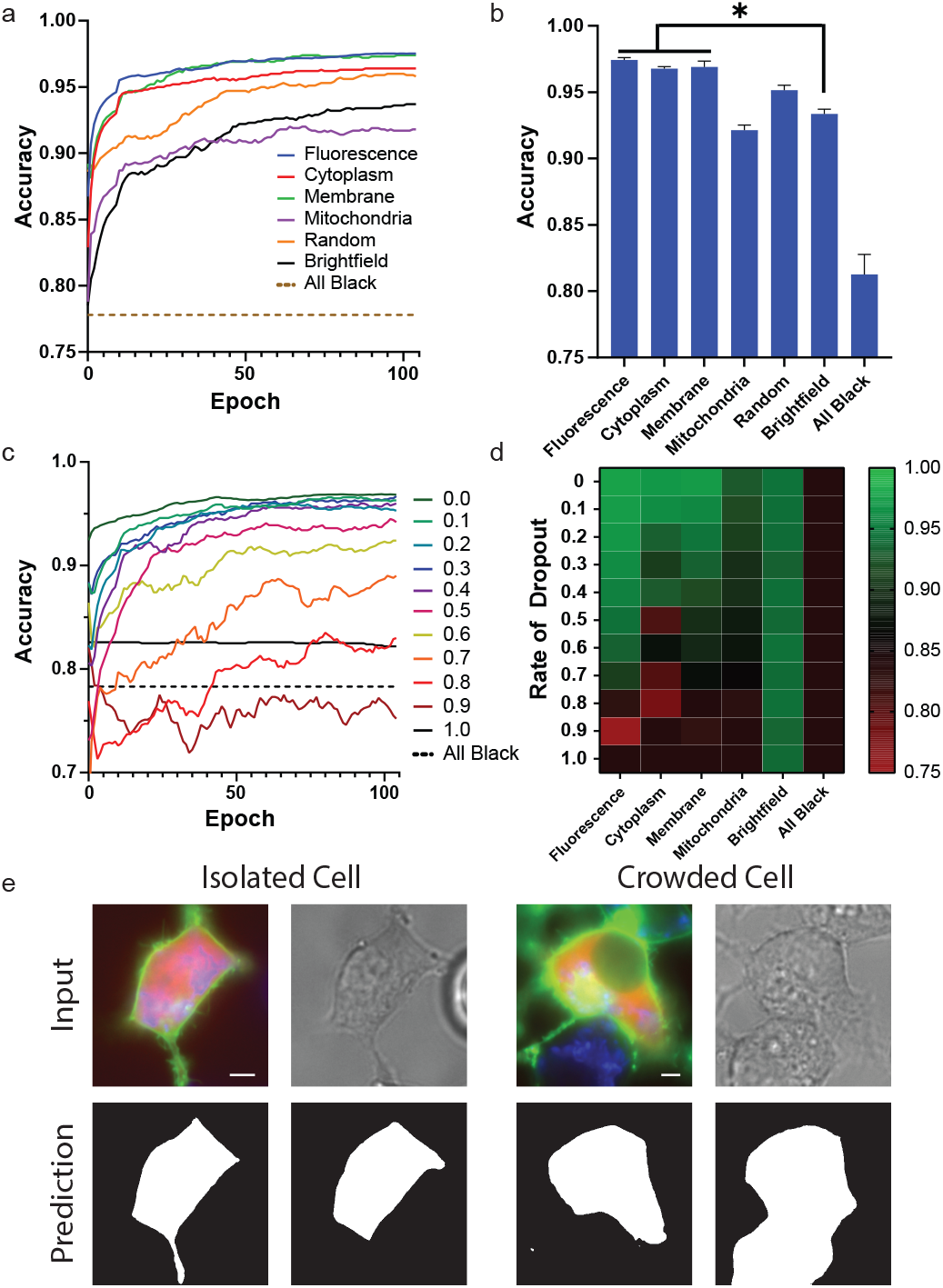
Comparison of segmentation accuracies when using common cell labeling approaches. (a) Representative training accuracies of models trained using subimages representing common segmentation techniques. (b) Final accuracies after training determined by a 10-point rolling window. Data are presented as mean ± s.e.m. \**P <* 0.05 compared to brightfield as determined by a paired t-test. All bars were significant compared to the “All Black” control. (c) Representative training accuracies of fluorescence-based models as the rate of fluorescence dropout was gradually increased. (d) Heatmap representing the final segmentation accuracy of models trained under various fluorescent dropout conditions. (e) Example segmentation results for a cell in a low or high density environment. Results are presented in the order of: input image and model prediction for both the fluorescence and brightfield subimages created from the same hyper-labeled source image. The images on the left represent a cell that is isolated from its neighbors while the cell on the right is in direct contact with neighboring cells. White scale bar represents 0.5 μm.

### B. Improving brightfield performance using out-of-focus (OoF) brightfield imaging

To determine why the brightfield model was particularly poor at distinguishing cell boundaries when cells were highly confluent, we first examined the intensity profiles. These data revealed cell boundaries are much more difficult to discern using brightfield images than their fluorescence counterparts, especially when cells are in close proximity (**Fig. 3a**). This mirrors human performance, where segmentation of confluent cells was much less accurate in brightfield images than in either cytoplasmic or membrane images (**Supplementary Fig S9**). We noticed in this profiling that cells that were slightly out-of-focus (OoF) in the brightfield image showed more stark diffraction patterns near the cell boundary. These diffraction patterns serve to either highlight or darken these edges, producing an intensity pattern with either peaks or valleys at the cell boundary (**Fig. 3a,b**). Supplementing the in-focus bright-field channel with one OoF image above the plane of focus (+3, +5, or +10 μm) and one below (−3, −5, or −10 μm) slightly improved performance compared to the in-focus brightfield model (**Fig. 3c,d**). However, it was found that the optimal focal distance was cell-type dependent (**Supplementary Fig. S6**), with thinner AD293 cells performing better at lower offsets and thicker INS1E cells performing best at higher offsets. To account for those differences, a segmentation model (RND Br A+B) was trained using subimage stacks composed of one of the lower OoF brightfield channels chosen at random, the infocus brightfield image, and one of the higher OoF brightfield channels chosen at random. Although this did not result in the best overall performance of the models tested, it performed reasonably well on both cell types (**Supplementary Fig. S6**) presenting a more robust approach to training cells of variable morphologies or heights. These data suggest that using OoF brightfield images provide a method of improving the baseline performance of label-free segmentation models, although care must be taken to account for the range of expected cell thicknesses and confluencies found across samples.

**Fig. 3.**
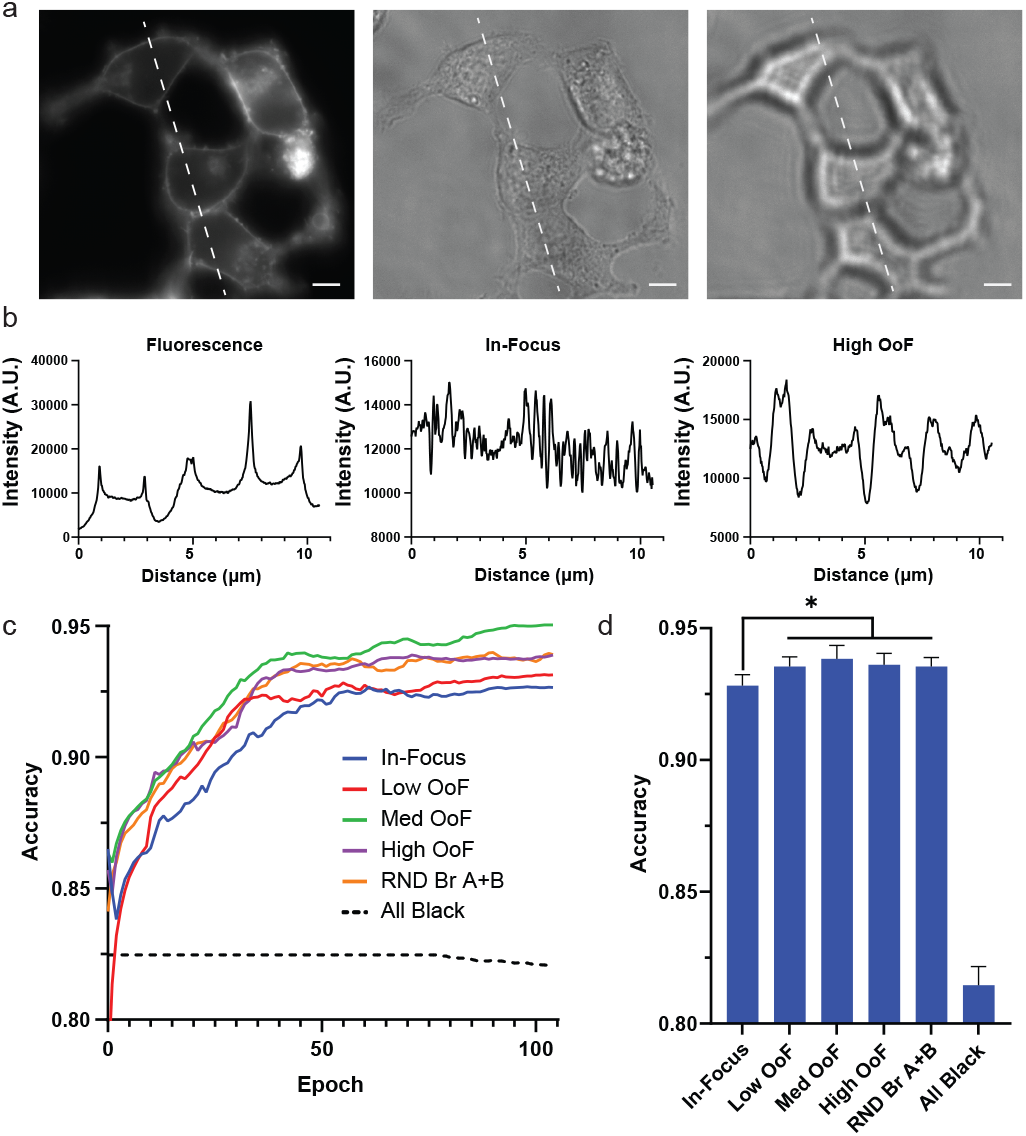
Analysis of brightfield imaging. (a) Representative images of cells expressing Membrane-tagged YFP imaged with fluorescence as well as a brightfield image taken in and out-of focus. White scale bar represents 1 μm. Lines drawn on the images represent the axis along which intensity plots (b) were taken. (c) Representative run of the models trained using various combinations of brightfield images. (d) Final accuracies of training (n=6). Data are presented as mean ± s.e.m. \**P <* 0.05 compared to in-focus brightfield alone as determined by a paired t-test.

### C. Fluorescence merging as a solution to uneven labeling

Despite the performance advantage afforded by incorporating OoF imaging, peak brightfield performance alone was still significantly below that of the fluorescence, cytoplasm, and membrane models (comparing **Fig. 2b** to **3d**). To combine the performance advantage of fluorescence with the reliability of brightfield under label-free conditions, we explored training segmentation models using combinations of fluorescence and brightfield channels (**Fig. 4**). For these models, subimages stacks were composed of one fluorescence channel, one infocus brightfield channel, and one OoF brightfield channel, with the fluorescence channel composed of either an individual channel or a merge of all the fluorescent channels available (see **Supplementary Fig. S2** and **Fig. 1d,e**). In all cases, the introduction of a fluorescence channel either maintained or improved the accuracy of brightfield alone (**Fig. 4b**), with the addition of a cytoplasmic, membrane, or merged fluorescence channel conferring the largest advantages. The performance of fluorescence-based models (cytoplasm, mitochondria, membrane, fluorescence) were previously found to substantially decrease under variable labeling conditions. To determine whether the inclusion of brightfield channels in the subimage stacks would guard against this effect, models were trained under a range of dropout rates as before. As the rate of dropout was increased, the fluorescence model dropped in performance significantly while the combination models each converged to the performance of the OoF brightfield model (RND Br A+B, **Fig. 4c**). These data suggest that training cell segmentation models using a combination of fluorescence and brightfield channels can produce a model that can effectively segment cells in the absence of fluorescence markers, while also being capable of capitalizing on fluorescent markers to improve performance when they are available. In particular, performing cell segmentation using subimage stacks composed of both a merged fluorescence and an OoF brightfield channel (the Merge + Br model) presents itself as a promising strategy to maximize performance across a range of labeling conditions.

**Fig. 4.**
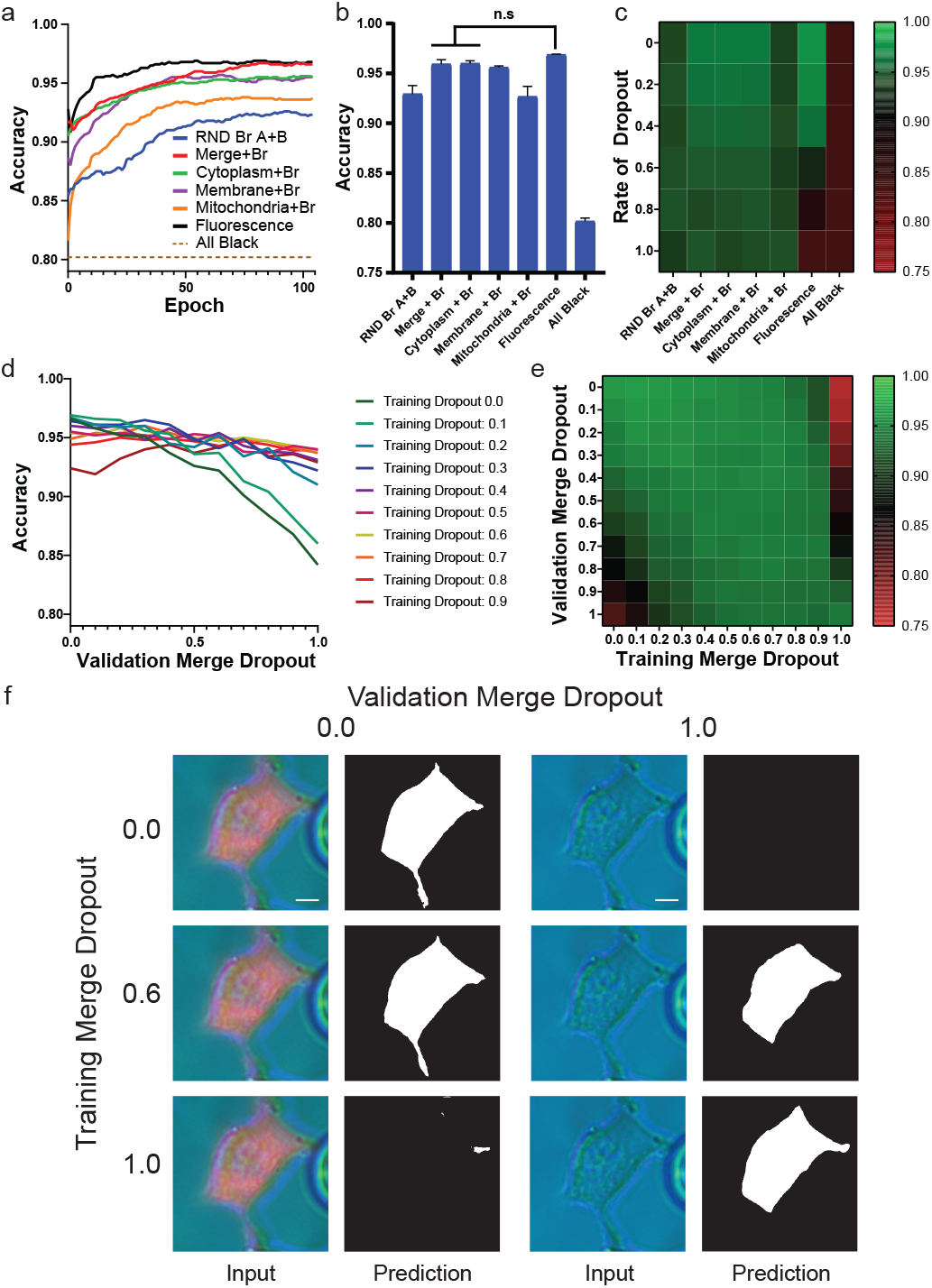
Combining fluorescence and brightfield techniques. (a) Representative run of models trained on various fluorescence/brightfield combinations. (b) Final accuracies of the training runs (n=3). Data are presented as mean ± s.e.m. \**P <* 0.05 compared to in-focus brightfield alone as determined by a paired t-test. (c) Heatmap of final training accuracies for each model as the rate of fluorescence dropout increased. Representative run (d) and pooled heatmap (e, n=3) relating validation accuracy for each model when exposed to data of varying merge dropout rates. (f) Representative image of a cell segmented under various training and validation conditions. Left image in each pair represents the merged image fed into the network; right image represents the resulting mask. White scale bar represents 0.5 μm.

Cell segmentation models are robust if they can maintain segmentation accuracy across a wide range of labeling conditions, which can be simulated by altering the rate of dropout during the validation set independent of the rate used to train that model. For instance, a model trained without any dropout (i.e. a training dropout rate of 0%) can then be independently validated using a dataset where either all fluorescent channels are replaced by black equivalents (i.e. a validation dropout rate of 100%), no channels are replaced (i.e. a validation dropout rate of 0%), or a mixture (1-99%). The robustness of a model can therefore be determined by its performance across the range of validation dropout rates spanning 0% to 100%. To determine how to maximize the robustness of the Merge+Br model, 11 models were trained using a single merge dropout rate (ranging from 0% to 100% in 10% increments). Each model was then validated across the full range of dropout rates (**Fig. 4d,e**). Although models trained using low dropout rates had the greatest overall accuracy on well-labeled datasets, they quickly lost accuracy as labeling became more sparse (**Fig. 4d**). Similarly, the model trained without fluorescence (i.e. a merge dropout rate of 100%) lost performance as labeling was introduced, indicating that the presence of novel information was disruptive to the model if not previously encountered. Instead, models trained using a merge dropout rate between 40-60% were the most consistent across the labeling spectrum (**Fig. 4e**). A closer examination of the segmentation outputs (**Fig. 4f**) from models trained using a dropout rate of 0%, 60% and 100% revealed that training a model using only one type of source input (i.e. 0% or 100%) obliterated its ability to segment cells when not presented with that same input. These data highlight the importance of maximizing the range of experimental conditions experienced by the model during training. These results were further validated using a test set (**Supplementary Fig. S7**).

To determine whether combining a fluorescence merge channel with brightfield images was generally beneficial to segmentation, we sourced an external dataset collected from a different cell type (macrophages) and differently stained with BODIPY 493/503 and Sytox to label the cytoplasm and nucleus, respectively [18]. The cells in this dataset were at significantly lower confluency (i.e. showed less cell-to-cell contact) and the brightfield images were collected using differential interference contrast (DIC), both of which limited the benefits of fluorescence and out-of-focus blur on edge recognition. Accordingly, segmentation using either fluorescence alone or the in-focus brightfield channel performed similarly (**Supplementary Fig. S8b**), and the inclusion of OoF brightfield images had a neutral effect (**Supplementary Fig. S8c**). However, the combination of brightfield and fluorescence merge resulted in increased segmentation accuracy as compared to either fluorescence or brightfield alone (**Supplementary Fig. S8d**). This increased segmentation accuracy was maintained in the face of inconsistent fluorescent labeling (**Supplementary Fig. S8e**). As with the primary dataset, training with a moderate dropout rate produced models that maintained their performance across a wider range of validation dropouts (**Supplementary Fig. S8f**). These data suggest that while each individual component may not contribute to increased segmentation performance for all applications (e.g. OoF brightfield), the combination of merged fluorescence and brightfield provides a robust approach to maximizing segmentation performance.

## IV Discussion

The unique nature of microscope images provides many exciting opportunities for innovation when adapting deep learning techniques for cell segmentation. Notable among them is the independence of channels within a microscope image stack, such that representative subimage stacks can be generated from a larger source, or hyper-labeled, image stack (**Fig. 1a**). Training segmentation models from these subimage stacks confers some key advantages, including the ability to (i) directly compare labeling approaches using identical cells (**Figs. 1b**; **2a,b**), (ii) keep dataset requirements small, and (iii) simulate experimental conditions during training (i.e. variable fluorescence labeling, see **Figs. 1e,f**; **2c,d**). Here, we demonstrate these advantages using a dataset comprised of image stacks constructed from three fluorescent tags (cytoplasmic, membrane, and mitochondrial) and seven brightfield images (each at different focal planes) to both compare the relative advantages of fluorescence and brightfield-based segmentation approaches, and explore novel strategies. Central to this comparison was the tradeoff between peak segmentation accuracy and consistency.

Fluorescence-based approaches boast strong accuracies for fully labeled cells (**Fig. 2a,b**), but performed poorly as labeling became increasingly sparse (**Fig. 2c,d**). In contrast, brightfield approaches had lower peak accuracy scores, but were label-independent. Improving general performance was therefore accomplished using a two-pronged approach: first, by improving the base performance of brightfield images and second, to make use of fluorescence information when available without relying on it explicitly. Improvements to brightfield performance were accomplished by augmenting brightfield images with out-of-focus (OoF brightfield) channels (**Fig. 3**), while fluorescence information was added through a bulk merge of all fluorescence signals (**Fig. 4**). What resulted was a model (named Merge+Br) that maintained performance across a wide range of labeling conditions (**Fig. 4a-c**). The Merge+Br approach to cell segmentation is particularly appealing as merging all available fluorescence channels renders both training and prediction label-agnostic. In other words, source images with differing numbers of available fluorescence channels can be processed identically during either model training or predictive purposes. These results also serve to highlight the utility of using subimage stacks generated from the larger source image stacks to minimize dataset size requirements and compare approaches. To account for all of the possible labeling combinations tested here without using subimage stacks would not only have required a prohibitively large dataset, but also lost the statistical power gained by comparison strategies on identical cells.

The ability to simulate various labeling conditions using subimage stacks was found to be particularly important when training models that are required to segment a more diverse set of experimental setups. For instance, we found that models trained on perfectly labeled cells boasted the highest accuracies but were also the quickest to lose performance as labeling became more inconsistent (**Fig. 4c**). In contrast, models trained on cells with varying degrees of fluorescence labeling (simulated using dropout rates between 40-60%) maintained a more consistent performance across the full range of labeling scenarios (**Fig. 4d,e**). Although training models in this manner may not produce the highest academic segmentation accuracies, it is critical for segmentation models used in image analysis pipelines where the set of available fluorescent labels may vary considerably across experiments.

A more subtle advantage of collecting hyper-labeled image stacks for training is that they expedite the creation of groundtruth labels. In certain cases, this may even allow ground-truth labeling to surpass human level performance. For instance, segmenting brightfield images of cells in close proximity is difficult for manual operators, but simple when a cytoplasmic or membrane marker is present (see **Supplementary Fig. S9**). Even if the final objective is to create a label-free segmentation model, imaging cells with a cytoplasmic or membrane marker in addition to the relevant brightfield channels allow more precise ground-truth labels to be created than through the brightfield channels alone. The label-free model can then be generated by training the model using subimage stacks containing only the relevant brightfield channels. By extension, combinations of markers may even be used in conjunction with other algorithmic or deep learning segmentation approaches to automate label generation for larger datasets, whether or not those specific markers will be used to train the model.

## ACKNOWLEDGMENT

This work was supported by the Natural Sciences and Engineering Research Council of Canada through Discovery (RGPIN-2016-371705) and Research Tool and Instrumentation (RTI-2018-00846) grants, and the Canada Foundation for Innovation Leaders Opportunity Fund. Stipend support was provided to W.D.C. by the Banting and Best Diabetes Centre (BBDC) and the Natural Sciences and Engineering Research Council of Canada (PGSD3-489652-2016), and to A.M.B. by an Ontario Graduate Scholarship (OGS 2018). We thank V. Pal and C. McFaul for technical assistance on the project.

## References

[1] Fuyong Xing and Lin Yang. Robust nucleus/cell detection and segmentation in digital pathology and microscopy images: a comprehensive review. IEEE reviews in biomedical engineering, 9:234–263, 2016.

[2] Alden A Dima, John T Elliott, James J Filliben, Michael Halter, Adele Peskin, Javier Bernal, Marcin Kociolek, Mary C Brady, Hai C Tang, and Anne L Plant. Comparison of segmentation algorithms for fluorescence microscopy images of cells. Cytometry Part A, 79(7):545–559, 2011.

[3] Olaf Ronneberger, Philipp Fischer, and Thomas Brox. U-net: Convolutional networks for biomedical image segmentation. In International Conference on Medical image computing and computer-assisted intervention, pages 234–241. Springer, 2015.

[4] Juan C Caicedo, Jonathan Roth, Allen Goodman, Tim Becker, Kyle W Karhohs, Matthieu Broisin, Molnar Csaba, Claire McQuin, Shantanu Singh, Fabian Theis, et al. Evaluation of deep learning strategies for nucleus segmentation in fluorescence images. BioRxiv, page 335216, 2019.

[5] Yousef Al-Kofahi, Alla Zaltsman, Robert Graves, Will Marshall, and Mirabela Rusu. A deep learning-based algorithm for 2-d cell segmentation in microscopy images. BMC bioinformatics, 19(1):365, 2018.

[6] Thorsten Falk, Dominic Mai, Robert Bensch, Özgün Çiçek, Ahmed Abdulkadir, Yassine Marrakchi, Anton BÖhm, Jan Deubner, Zoe Jäckel, Katharina Seiwald, et al. U-net: deep learning for cell counting, detection, and morphometry. Nature methods, 16(1):67, 2019.

[7] Shan E Ahmed Raza, Linda Cheung, David Epstein, Stella Pelengaris, Michael Khan, and Nasir M Rajpoot. Mimo-net: A multi-input multioutput convolutional neural network for cell segmentation in fluorescence microscopy images. In 2017 IEEE 14th International Symposium on Biomedical Imaging (ISBI 2017), pages 337–340. IEEE, 2017.

[8] Kaiming He, Xiangyu Zhang, Shaoqing Ren, and Jian Sun. Delving deep into rectifiers: Surpassing human-level performance on imagenet classification. In Proceedings of the IEEE international conference on computer vision, pages 1026–1034, 2015.

[9] Nitish Srivastava, Geoffrey Hinton, Alex Krizhevsky, Ilya Sutskever, and Ruslan Salakhutdinov. Dropout: a simple way to prevent neural networks from overfitting. The journal of machine learning research, 15(1):1929–1958, 2014.

[10] Diederik P Kingma and Jimmy Ba. Adam: A method for stochastic optimization. arXiv preprint arXiv:1412.6980, 2014.

[11] Leslie N Smith. A disciplined approach to neural network hyper-parameters: Part 1–learning rate, batch size, momentum, and weight decay. arXiv preprint arXiv:1803.09820, 2018.

[12] Leslie N Smith. Review of advanced imaging techniques Journal of pathology informatics, 3, 2012.

[13] Caicedo, Juan C and Cooper, Sam and Heigwer, Florian and Warchal, Scott and Qiu, Peng and Molnar, Csaba and Vasilevich, Aliaksei S and Barry, Joseph D and Bansal, Harmanjit Singh and Kraus, Oren Data-analysis strategies for image-based cell profiling. Nature Methods, 14(9):849, 2017.

[14] Ellen C Jensen. Use of fluorescent probes: their effect on cell biology and limitations. The Anatomical Record: Advances in Integrative Anatomy and Evolutionary Biology, 295(12):2031–2036, 2012.

[15] Ram Dixit and Richard Cyr. Cell damage and reactive oxygen species production induced by fluorescence microscopy: effect on mitosis and guidelines for non-invasive fluorescence microscopy. The Plant Journal, 36(2):280–290, 2003.

[16] Raphael Alford, Haley M Simpson, Josh Duberman, G Craig Hill, Mikako Ogawa, Celeste Regino, Hisataka Kobayashi, and Peter L Choyke. Toxicity of organic fluorophores used in molecular imaging: literature review. Molecular imaging, 8(6):7290–2009, 2009.

[17] William D Cameron, Cindy V Bui, Ashley Hutchinson, Peter Loppnau, Susanne Gräslund, and Jonathan V Rocheleau. Apollo-nadp+: a spectrally tunable family of genetically encoded sensors for nadp+. Nature methods, 13(4):352, 2016.

[18] Selinummi, Jyrki and Ruusuvuori, Pekka and Podolsky, Irina and Ozinsky, Adrian and Gold, Elizabeth and Yli-Harja, Olli and Aderem, Alan and Shmulevich, Ilya. Bright field microscopy as an alternative to whole cell fluorescence in automated analysis of macrophage images. PloS one, 4(10):e7497, 2009.

[19] Kaiming He, Xiangyu Zhang, Shaoqing Ren, and Jian Sun. Deep residual learning for image recognition. In Proceedings of the IEEE conference on computer vision and pattern recognition, pages 770–778, 2016.

[20] Yusuke Sugawara, Sayaka Shiota, and Hitoshi Kiya. Super-resolution using convolutional neural networks without any checkerboard artifacts. In 2018 25th IEEE International Conference on Image Processing (ICIP), pages 66–70. IEEE, 2018.

[21] Jeremy Howard et al. FastAI. https://github.com/fastai/fastai, 2018.

