## Supplemental figures for "Cell segmentation using deep learning: comparing label and label-free approaches using hyper-labeled image stacks"

### APPENDIX

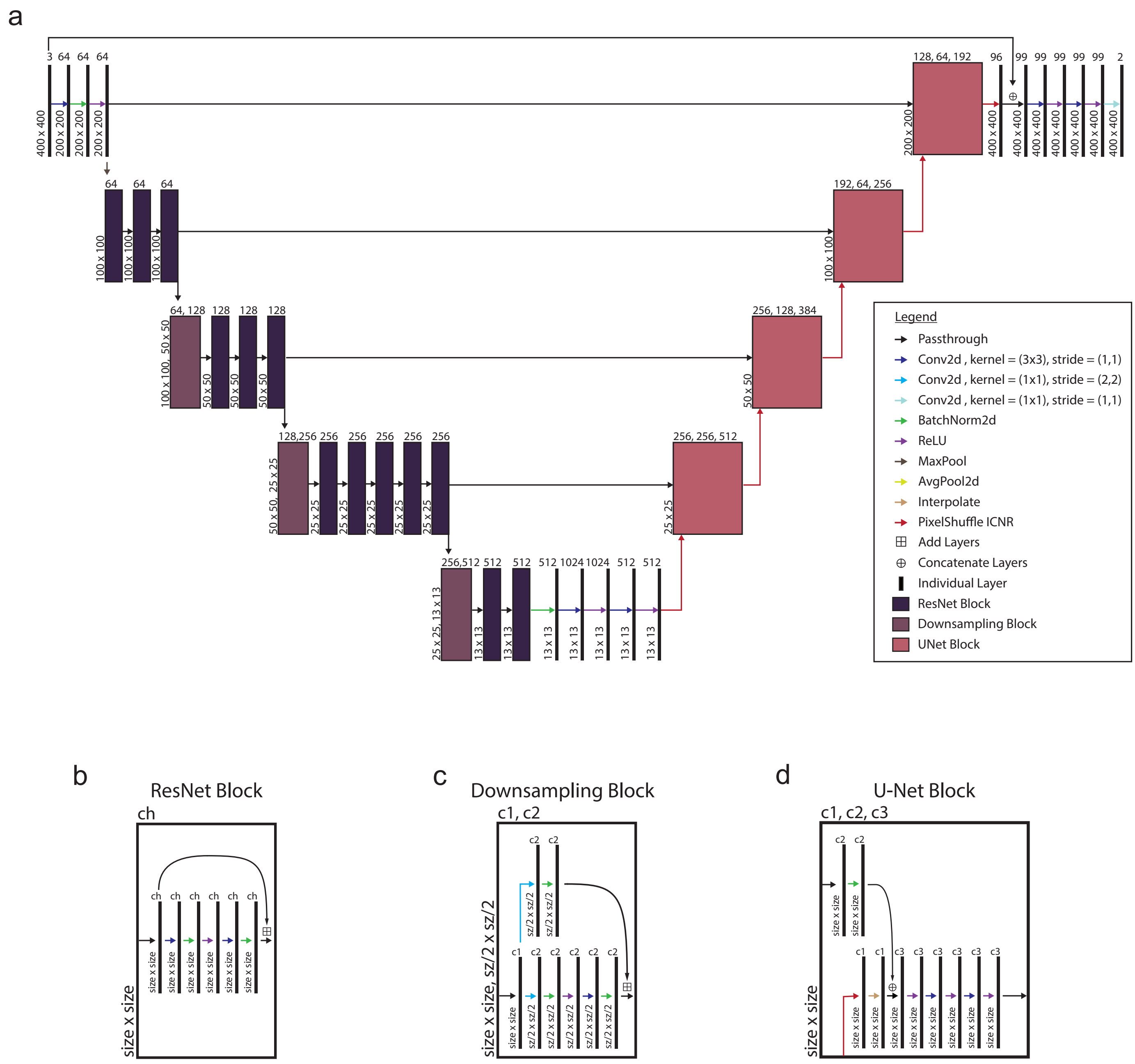


Fig. S1. Architecture of the custom U-Net architecture used to train the segmentation models. (a) Schematic of the architecture. Lines represent individual layers while rectangular blocks represent the common sequences of layers outlined in (b-c). Black arrows represent a direct passthrough of data while colored arrows represent that the indicated operation was executed on the input to the next layer. For an image of initial size 400 x 400 px, the numbers at the bottom left of each layer represent the image height and width, while the number on top represents the number of channels. (b) The operations involved in the ResNet block, which takes a single input and produces a single output. (c) The downsampling block, which receives a single input and produces a single output. sz/2 represents $\lceil size/2 \rceil$. (d) U-Net block, which takes two inputs and produces a single output.

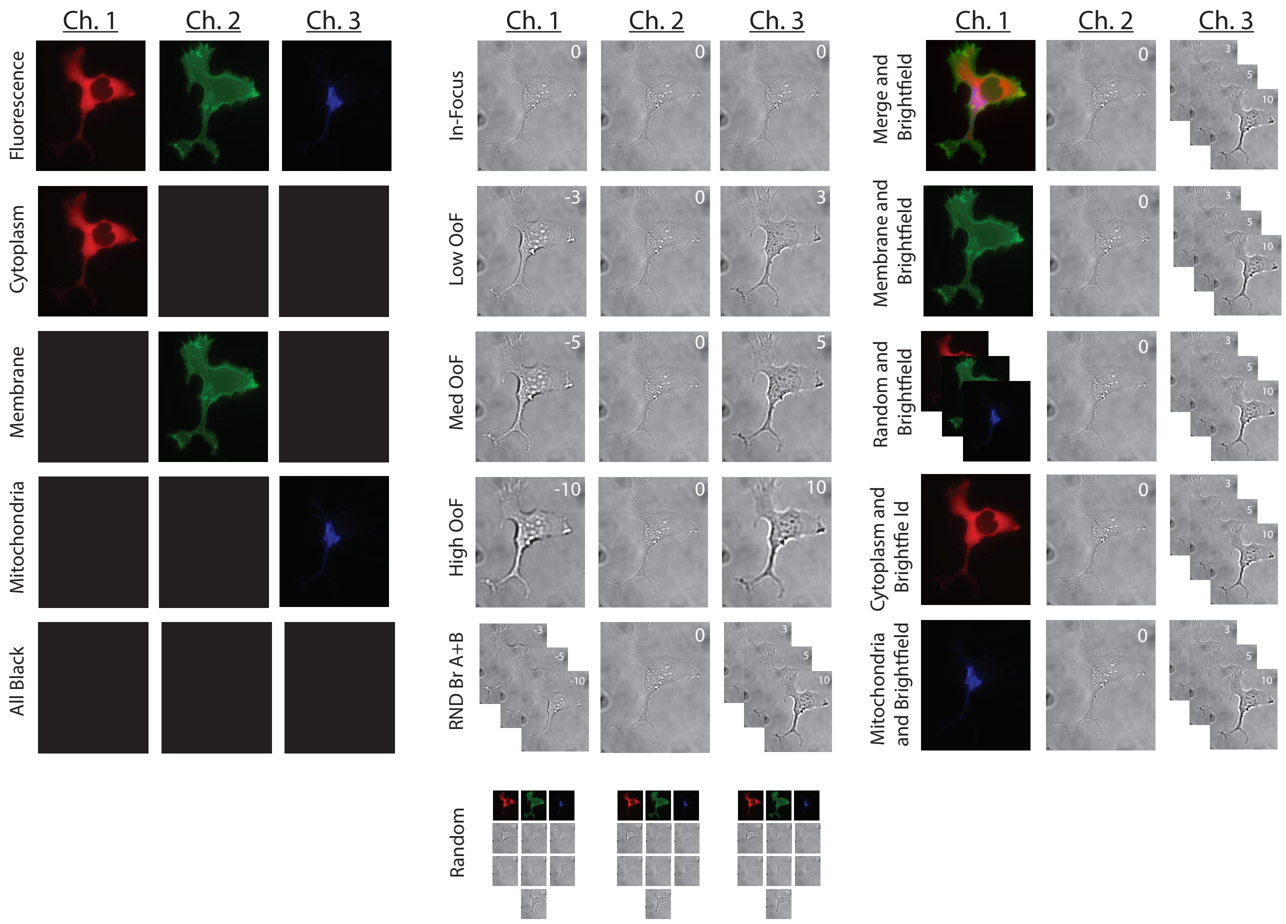


Fig. S2. Examples of subimage combinations used during training. Each model accepts an input of three channels according to a preprogrammed loading code. The presence of multiple images in the same channel slot (ex. "RND Br A+B") indicates that one of these channels was chosen at random with equal probability. For the first channel in "Merge and Brightfield", the three fluorescence channels were combined into a single channel. Colorization of the subchannels is used here for visualization purposes and does not represent multiple RGB channels. For brightfield channels, the number in the upper right corner represents the focal plane in μm. OoF represents the initialism of "Out-of-Focus".

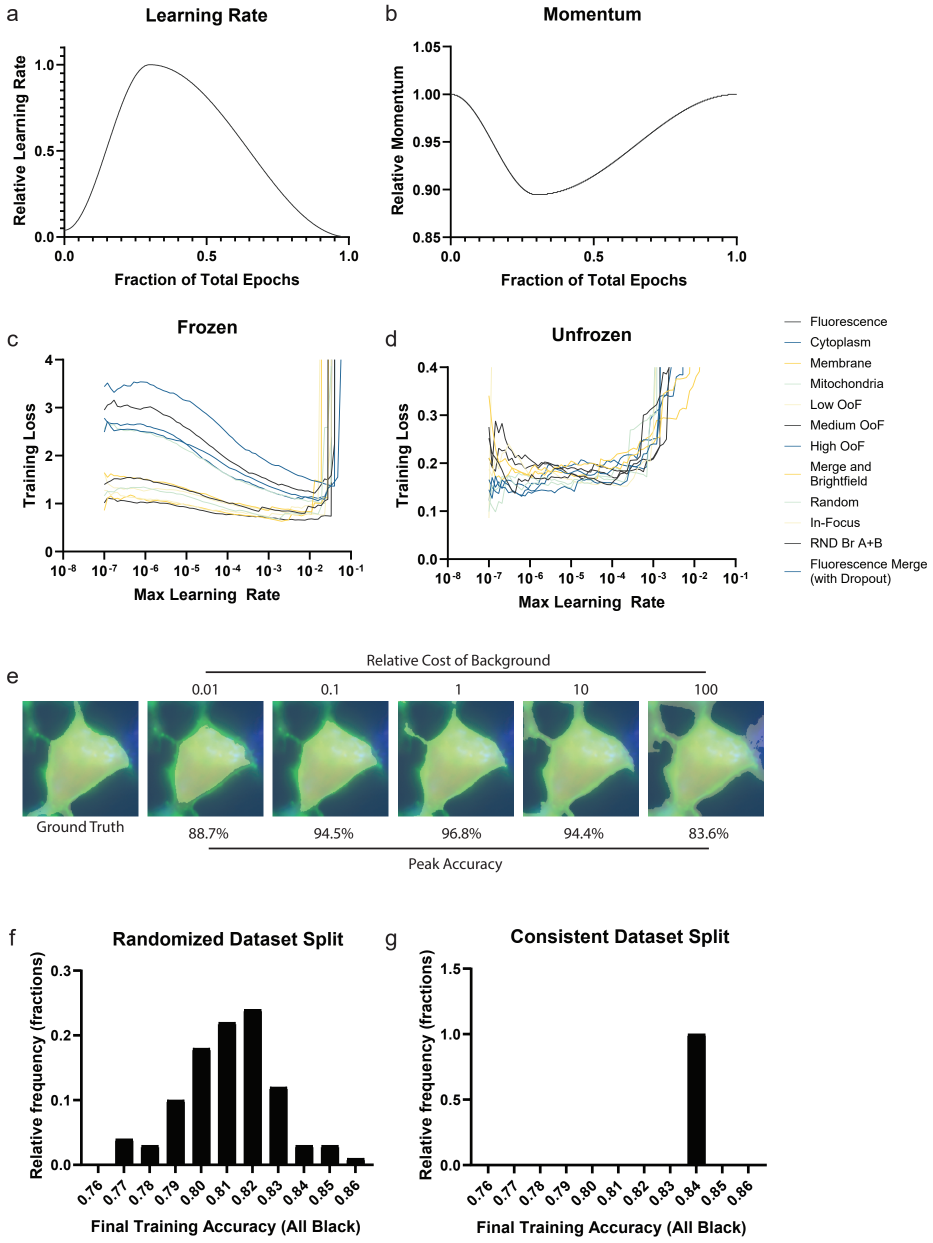


Fig. S3. Optimization of training parameters. (a-b) Learning rate and momentum were inversely scheduled based on a maximum learning rate. The relationship between training loss and the maximum learning rate was measured when the pre-trained descending arc was either (c) frozen or (d) unfrozen and used to determine the maximum learning rates for training. (e) Segmentation performance using weighted cross-entropy loss was compared across various weights for background loss. Segmentation accuracy using all black channels when training using a different 80:20 randomized training-validation split for each run (f) vs. the same training-validation split(g). Each graph represents 100 trials. Data from (f) was confirmed to be normally distributed through Shapiro-Wilk.

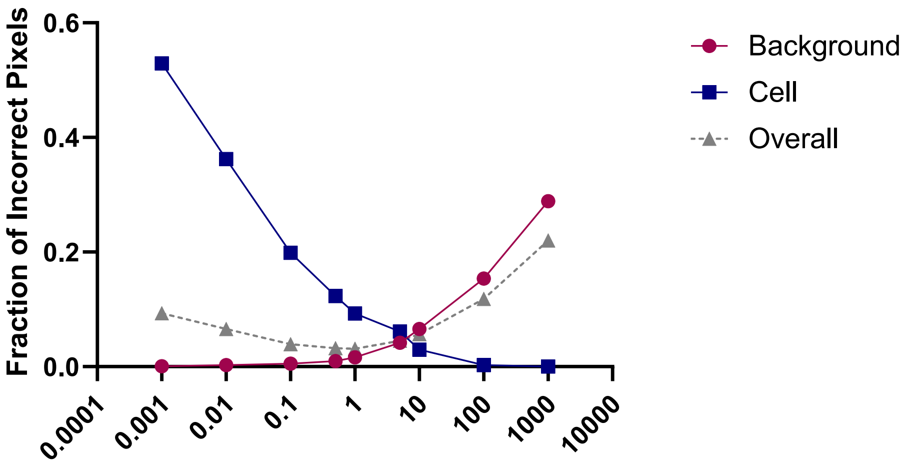


Fig. S4. Effect of cross entropy weighting on pixel accuracy. Models were trained using cross-entropy loss with a range of weights for the background class. The fraction of incorrect pixels was calculated by taking the average number of misclassified pixels from each image for the background class (red circles), target cell (black squares), or combination of both classes (grey triangles).

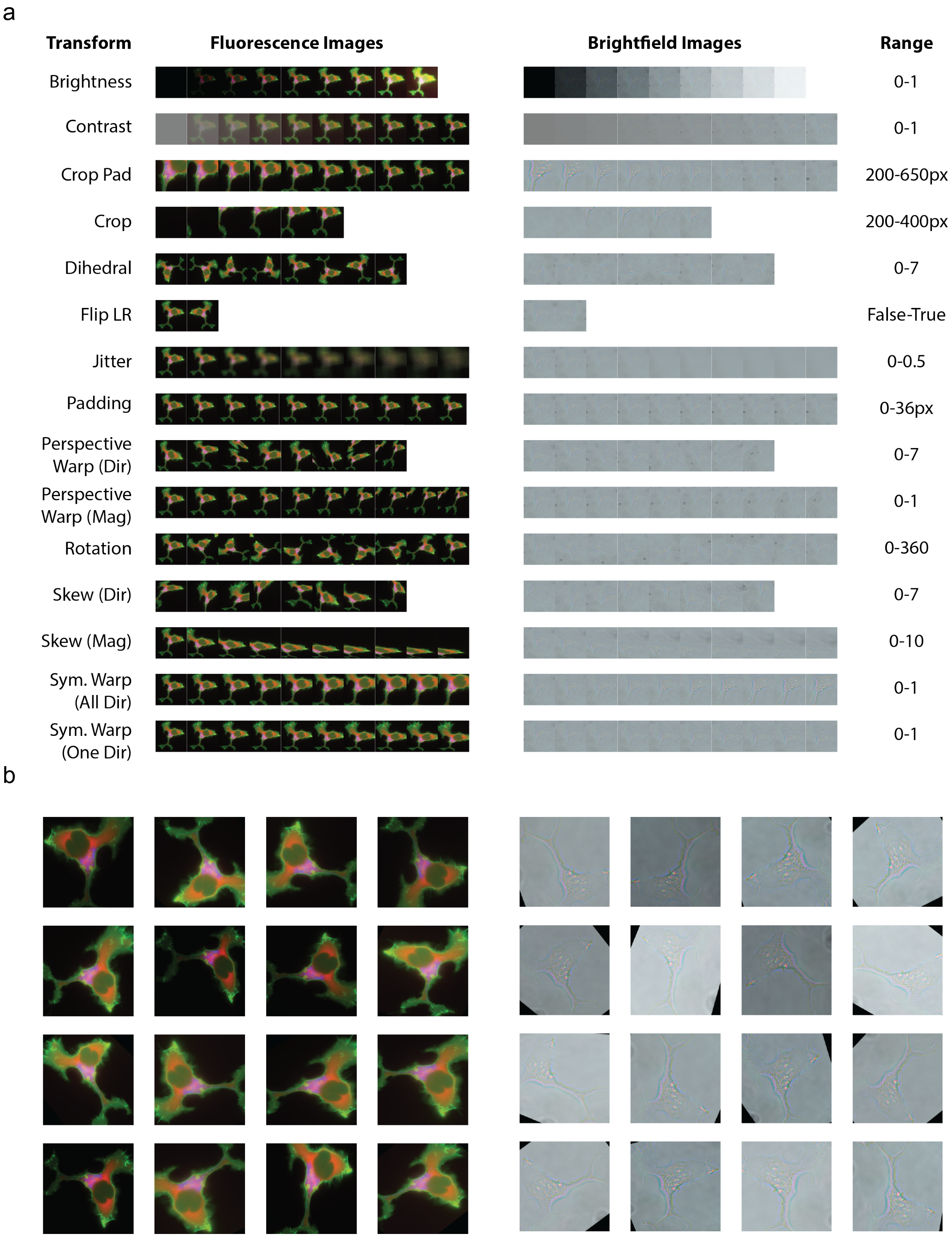


Fig. S5. Exploration of image transformations. (a) Individual transforms applied using a range of values to a representative brightfield and fluorescence image. "Skew" and "perspective warp" require both a direction and magnitude thus were investigated independently. For "skew" directions, the magnitude was 2; for "perspective warp" directions, the magnitude was 1. The "crop pad" transformation used border padding while reflection padding was used for all other transforms when needed. (b) Application of the chosen transforms to each image 16 times to demonstrate the expected range of inputs during training.

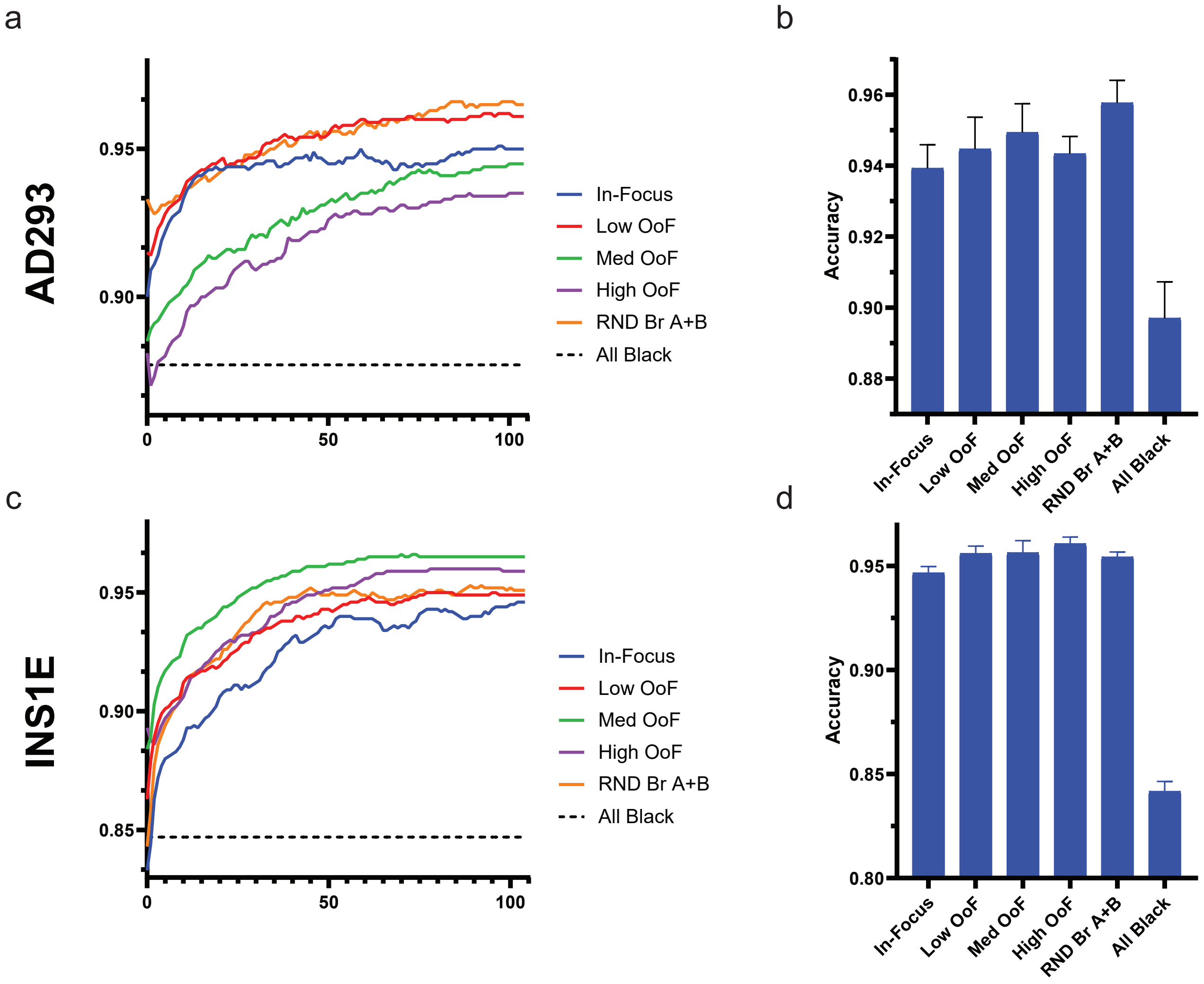


Fig. S6. Comparison of Out-of-Focus imaging on different cell types. Representative training curve for Out-of-Focus (OoF) models for (a) AD293 or (c) INS1E cells as well as their pooled final accuracies (b, d respectively). Pooled accuracies represent n=3.

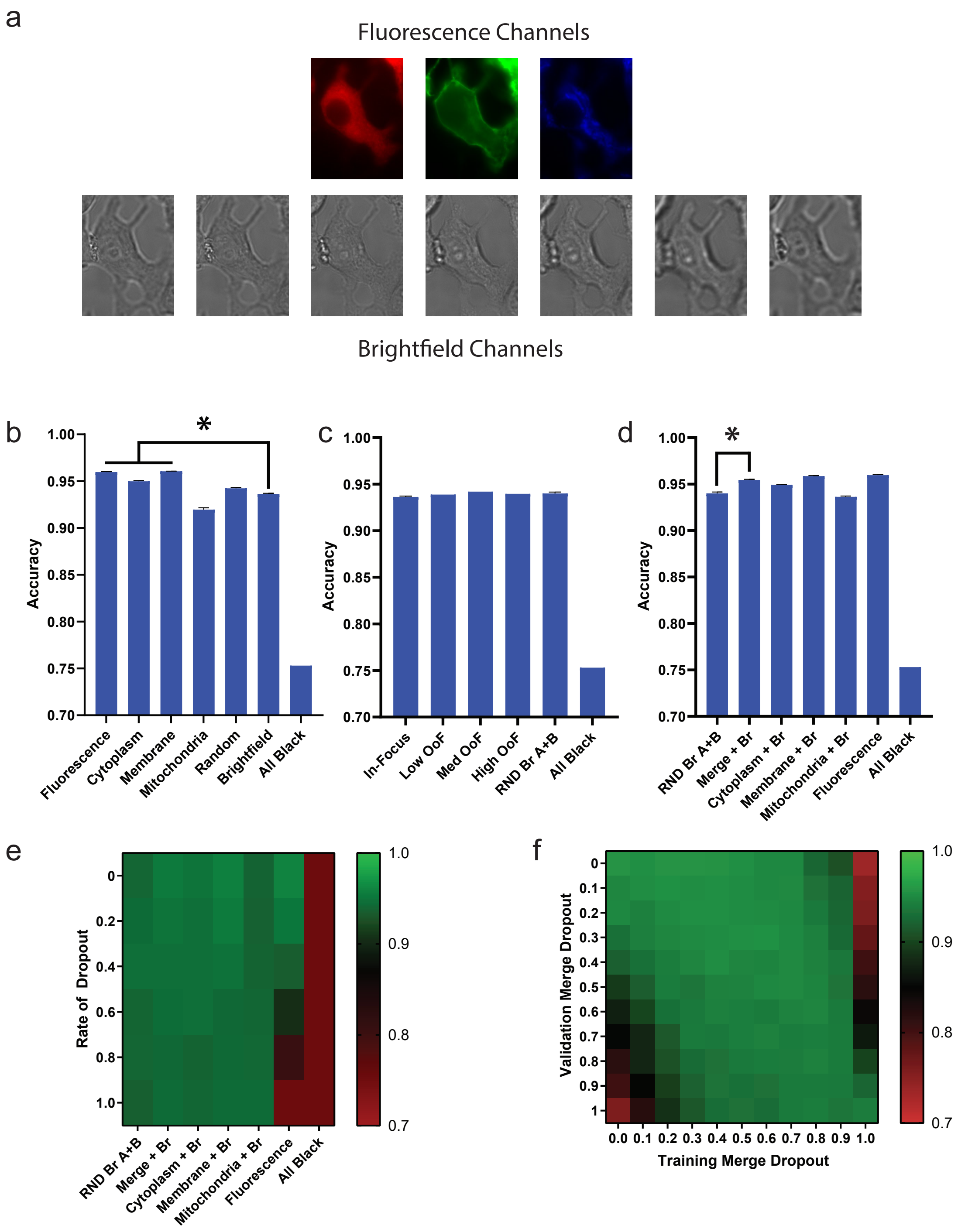


Fig. S7. Validation of main results using a test set. (a) Representative image from the test set. Cells were labeled with three distinct subcellular markers for: the ER (mTurquoise2-tagged Apollo-NADP$^{+}$), the membrane (YFP-Mem), and the mitochondrial matrix (Mitotracker Deep-Red FM). Brightfield images were taken of each cell at 7 different focal depths (-10, -5, -3, 0, +3, +5, +10 µm), with 0 µm representing cells in manual focus. (b-f) Segmentation accuracies were assessed by validating the external dataset on the models trained using the training dataset (discussed in section II-A). The graphs assess earlier results: (b) comparison of common labeling approaches (see Fig. 2b), (c) comparison of OoF imaging (see Fig. 3c), (d) comparison of mixed fluorescence and brightfield (see Fig. 4b), (e) model performance under dropout (see Fig. 4c), and (f) effect of training dropout rate on accuracy in face of a range of validation dropouts (see Fig. 4e). Data are presented as mean ± s.e.m. $^{*}P < 0.05$ as determined by a paired t-test.

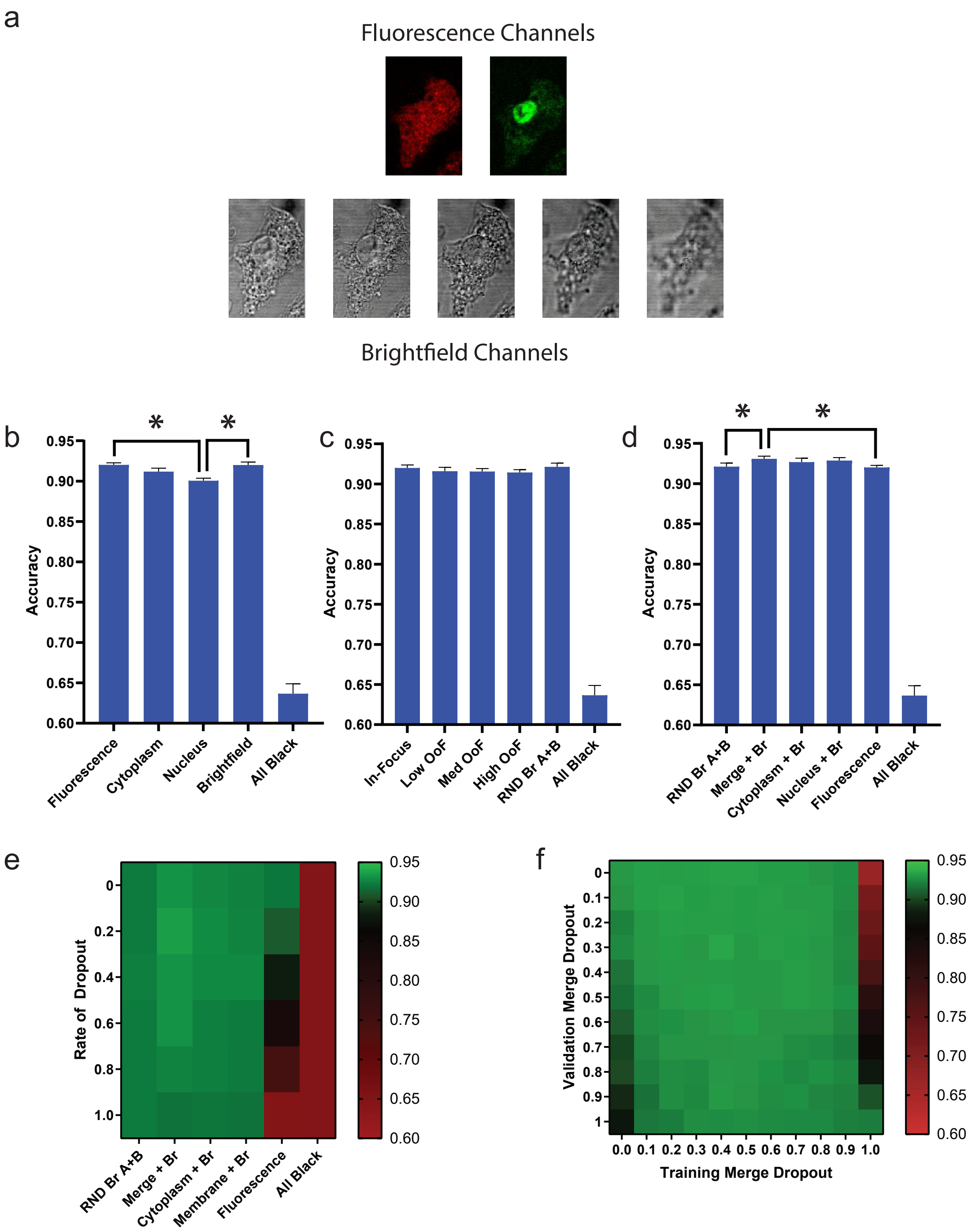


Fig. S8. Validation of methodology on an external dataset. External dataset consists of macrophage cells unstimulated and stimulated with lipopolysaccharide. (a) Representative image from the external dataset. Cells were labeled with two distinct subcellular markers for: the cytoplasm (BODIPY 493/503) and the nucleus (Sytox). Brightfield images were taken of each cell at 5 different focal depths. (b-f) Segmentation accuracies were assessed by validating the external dataset on the models trained using the training dataset (discussed in section II-A). The graphs assess earlier results: (b) comparison of common labeling approaches (see Fig. 2b), (c) comparison of OoF imaging (see Fig. 3c), (d) comparison of mixed fluorescence and brightfield (see Fig. 4b), (e) model performance under dropout (see Fig. 4c), and (f) effect of training dropout rate on accuracy in face of a range of validation dropouts (see Fig. 4e). Data are presented as mean ± s.e.m. $*P < 0.05$ as determined by a paired t-test.

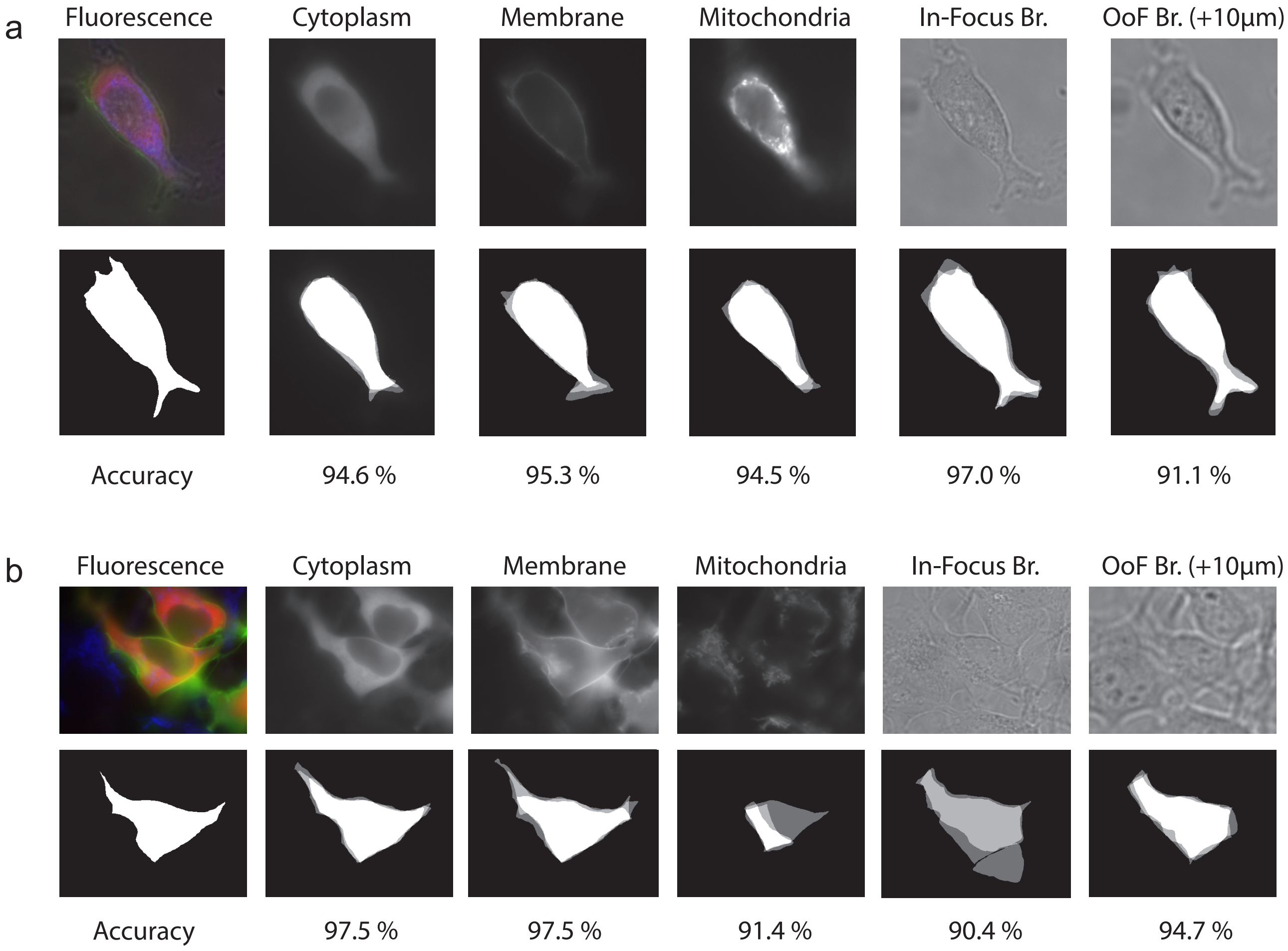


Fig. S9. Assessment of manual segmentation. Segmentation results of individual operators when segmenting using only one channel from the hyper-labeled image on cell in (a) isolation, or (b) confluency. Segmentation maps represent the average of 3 operators and accuracy was assessed as the percentage of pixels correctly identified, using the formula $(1 - y_{pred})(1 - y) + (y_{pred})(y)$ to assess individual pixel accuracy.
